# Cas9/CRISPR genome editing to demonstrate the contribution of Cyp51A Gly138Ser to azole resistance in *Aspergillus fumigatus*

**DOI:** 10.1101/311712

**Authors:** Takashi Umeyama, Yuta Hayashi, Hisaki Shimosaka, Tatsuya Inukai, Satoshi Yamagoe, Shogo Takatsuka, Yasutaka Hoshino, Minoru Nagi, Shigeki Nakamura, Katsuhiko Kamei, Kenji Ogawa, Yoshitsugu Miyazaki

## Abstract

Azole resistance in *Aspergillus fumigatus* is predominantly associated with increased expression of Cyp51A (lanosterol 14α-demethylase), the target enzyme of azole antifungal agents, or with single-nucleotide polymorphisms (SNPs) in *cyp51A*. Although several SNPs that may be linked to low susceptibility in azole-resistant isolates have previously been reported, few studies have been conducted to conclusively demonstrate the contribution of SNPs to decreased azole susceptibility. An *A. fumigatus* strain was isolated from the sputum of a 74-year-old male receiving long-term voriconazole treatment for chronic progressive pulmonary aspergillosis. Etest antifungal susceptibility testing showed low susceptibility to voriconazole, itraconazole, and posaconazole. Nucleotide sequencing of *cyp51A* from this isolate revealed the mutations Gly138Ser (GGC→AGC) and Asn248Lys (AAT→AAA) compared with the *cyp51A* of azole-susceptible isolates. PCR-amplified DNA fragments containing *cyp51A* with or without the mutations of interest and a hygromycin marker were simultaneously introduced along with the Cas9 protein and *in vitro*-synthesized single-guide RNA into protoplasts of the azole-resistant/susceptible strains. Etest azole susceptibility testing of recombinant strains showed an increased susceptibility via the replacement of Ser138 by glycine. In contrast, azole susceptibility was slightly decreased when a Ser138 mutation was introduced into the azole-susceptible strain AfS35, indicating that the serine at position 138 of Cyp51A contributes to low susceptibility in the azole-resistant isolate. Genetic recombination, which has been hampered thus far in clinical isolates, can now be achieved using Cas9/CRISPR genome editing. This technique could be useful to investigate the contribution of other SNPs of *cyp51A* to azole resistance.

## Introduction

The filamentous fungi *Aspergillus fumigatus* is the most common opportunistic human fungal pathogen, with a wide range of clinical features including invasive pulmonary aspergillosis, chronic progressive pulmonary aspergillosis (CPPA), and allergic bronchopulmonary aspergillosis (1). Triazole antifungal drugs are the most common treatment for *A. fumigatus* infection. Itraconazole (ITC) and voriconazole (VRC) are the only oral drug treatment options for aspergillosis, which may lead to long-term administration. Since the discovery of the first ITC-resistant isolate in 1997 (2), epidemiological reports of new triazole-resistant isolates have been increasing worldwide (3). Mechanisms of acquired azole resistance may be explained by extended periods of azole exposure in the host or by environmental exposure of *A. fumigatus* to agricultural fungicides.

The primary molecular mechanisms of triazole resistance in *A. fumigatus* isolates are mutations that alter the target protein Cyp51A and prevent its interaction with the drug (4). Mutations in *cyp51A* may be classified as single-nucleotide polymorphisms (SNPs) and/or tandem repeats in the promoter region (3). The major SNPs affecting Cyp51A are positioned at Gly54, Gly138, Met220, and Gly448. The clinical isolates having these SNPs demonstrate various azole susceptibility profiles; for example, isolates with SNPs at Gly54 show resistance to ITC and varied susceptibility to posaconazole (POS) and VRC, whereas isolates with SNPs at Gly138 show resistance to pan-azoles, including ITC, POS, and VRC. Another alteration is a tandem repeat in the promoter region that results in the overexpression of *cyp51A*. Two major classes of such azole-resistant mutants are TR34/Leu98His and TR46/Tyr121Phe/Thr289Ala, which carry a 34-bp and a 46-bp sequence duplication, respectively, as well as amino acid substitutions.

Although many SNPs in *cyp51A* that may be linked to low susceptibility in azole-resistant isolates have been previously reported, few studies have been conducted to conclusively demonstrate the contribution of SNPs to decreased azole susceptibility in clinical isolates. One obstacle affecting the molecular analysis of clinical *A. fumigatus* isolates is the production of genetically manipulated mutants, as the efficiency of homologous recombination is extremely low.

Cas9/CRISPR (the clustered regularly interspaced short palindromic repeats) is essentially a bacterial defense system for adaptive immunity against invading nucleic acids and has been applied as a powerful genome editing tool in various organisms (5). By forming a ribonucleoprotein complex with an artificial single-guide RNAs (sgRNAs) designed to target a cellular gene, the Cas9 nuclease efficiently introduces double-stranded breaks (DSBs) at the corresponding target locus (6). The sgRNA hybridizes to its complementary DNA sequence, immediately upstream of the protospacer adjacent motif (PAM), which consists of NGG for the *Streptococcuspyrogenes* Cas9 variant (7). DSBs in the target genomic DNA can be repaired either by homology-directed repair or non-homologous end joining (NHEJ) (5, 8, 9). DNA repair via homology-directed repair requires a homologous DNA template with sequence similarity to the adjacent region of the DSB locus, whereas NHEJ ligates the DSB, leading to indels in a template-independent manner. The Cas9/CRISPR system has also been successfully applied to *A. fumigatus* (10–12).

In the present study, we investigated the antifungal mechanisms of a pan-azole-resistant strain isolated from a patient receiving long-term VRC treatment for CPPA. Although two polymorphisms were found in *cyp51A* from the isolate, it was unclear which of the SNPs contributed to low azole susceptibility. We genetically demonstrated that one of the SNPs predominantly contributes to low susceptibility in the azole-resistant clinical isolate using a Cas9/CRISPR genome editing technique.

## Methods

### *A. fumigatus* strains and media

The *A. fumigatus* strains used in the present study are listed in Table 1. A clinical isolate NIID0345 was obtained in 2016 from the sputum sample of a 74-year-old male patient with CPPA, who had received VRC treatment for 3 years. *A. fumigatus* cultures were routinely grown in *Aspergillus* minimal medium (AMM: 10 g glucose, 0.516 g KCl, 0.516 g MgSO_4_·7H_2_O, 1.516 g KH_2_PO_4_, 1.516 g Mg(NO_3_)2·6H_2_O, 1 mL trace elements (13) in 1 L distilled water), Czapek-Dox medium (CD, BD Difco Laboratories Inc., Franklin Lakes, NJ), YG medium (13), or potato dextrose agar medium (PDA, BD Difco). For solid medium, 1.5% agar was added. *A. fumigatus* conidia were obtained from mycelia cultured on AMM or PDA at 30°C for 3–7 days, harvested with PBS containing 0.05% (v/v) Tween 20 and 20% (v/v) glycerol, and filtered through a 40-μm nylon cell strainer (Greiner Bio-One, Germany).

**Table 1.** *Aspergillus fumigatus* strains used in this study.

| Strain | Parent | Genotype<sup>a</sup> | Source |
| --- | --- | --- | --- |
| NIID0345 | clinical isolate |  | Current study |
| NIID0345-mut1-2 | NIID0345 | mut1 mut2 *cyp51A hph* | Current study |
| NIID0345-S138G | NIID0345 | mut1 S138G mut2 *cyp51A hph* | Current study |
| NIID0345-K248N | NIID0345 | mut1 K248N mut2 *cyp51A hph* | Current study |
| NIID0345-S138G-K248N | NIID0345 | mut1 mut2 *cyp51A hph* | Current study |
| AfS35 | D141 | *akuAΔloxP* | Fungal genetic<br>stock center |
| AfS35-mut1-2 | AfS35 | mut1 mut2 *cyp51A hph* | Current study |
| AfS35-G138S | AfS35 | mut1 G138S mut2 *cyp51A hph* | Current study |

<sup>a</sup> mut1 and mut2 are silent mutations for Cas9-nuclease resistance.

### DNA extraction, PCR, and sequencing

Genomic DNA extractions and purifications were performed using a DNeasy Plant Mini Kit (QIAGEN, Germany). Primers for the amplification and sequencing of *cyp51A* are listed in Table 2. Identification was confirmed by sequencing of the internal transcribed spacer (ITS) and D1/D2 regions and the β-tubulin gene. PCR amplification of *cyp51A* was performed using NIID0345 genomic DNA as a template and primers Discheck5 and Discheck3 using Q5 Hot Start High-Fidelity 2× Master Mix (New England Biolabs, Ipswich, MA).

**Table 2.**
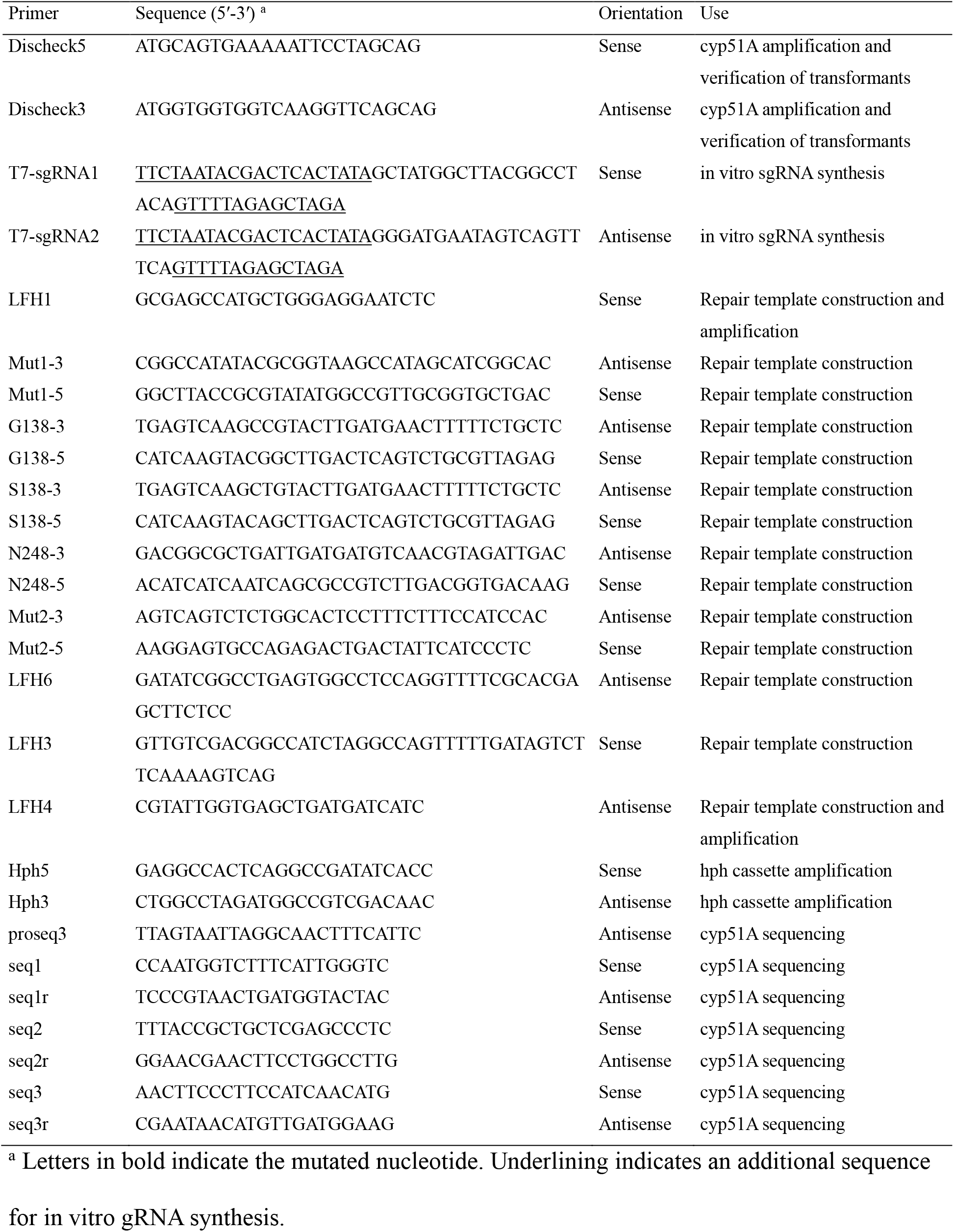
Oligonucleotide primers used in this study.

### sgRNA *in vitro* synthesis

We manually searched for target sequences consisting of G(N)15(A/T)(N)3NGG near the N-terminus (for sgRNA1) and C-terminus (for sgRNA2) as sgRNA target sequences and synthesized two oligonucleotides (T7-sgRNA1 and T7-sgRNA2, Table 2) consisting of the T7 promoter, sgRNA target sequence, and overlap sequence with Cas9 scaffold. These oligonucleotides were used for sgRNA synthesis via the EnGen® sgRNA Synthesis Kit, *S. pyogenes* (New England Biolabs). The synthesized sgRNAs were purified using an RNA clean & concentrator-25 (Zymo Research, Irvine, CA), quantified using a QuantiFluor RNA system (Promega, Madison, WI) and Quantus Fluorometer (Promega), and used for ribonucleoprotein formation with Cas9.

### Repair templates

A pHph plasmid harboring a hygromycin B resistance cassette (*hph*) was generated by deletion of two *loxP* sequences and HSV1 thymidine kinase sequences from pSK397 (14). Primers for the repair template construction are listed in Table 2. A region from 825-bp upstream to 1503-bp downstream of the *cyp51A* coding region was used for repair templates. The mutations and *hph* marker were introduced via PCR sewing or overlap extension PCR. The *hph* marker for selection of transformants was inserted between nucleotides 500 and 501 downstream of the *cyp51A* stop codon. Q5 Hot Start High-Fidelity 2× Master Mix (New England Biolabs) was used for PCR amplification. Primer combinations for overlap extension PCR are listed in Table 3. Briefly, NIID0345 or AfS35 genomic DNA was used as a template to generate overlapping PCR products with the corresponding site-specific mutations or junctions between *cyp51A* and the *hph* marker. The overlapping PCR products were mixed together and used as a template in the PCR-sewing step using the primers LFH1 and LFH4. Overlapping PCR product combinations are listed in Table 4. The fused PCR products were purified using a NucleoSpin® Gel and PCR Clean-up kit (Takarabio, Japan) and used for *A. fumigatus* protoplast transformation.

**Table 3.** Combination of primers for overlapping PCR used in this study.

| Name of PCR product | Primers | Template DNA |
| --- | --- | --- |
| 0345-A | LFH1/Mut1-3 | NIID0345 genomic DNA |
| 0345-mut1-G138 | Mut1-5/G138-3 | NIID0345 genomic DNA |
| 0345-mut1-N248 | Mut1-5/N248-3 | NIID0345 genomic DNA |
| 0345-mut1-mut2 | Mut1-5/Mut2-3 | NIID0345 genomic DNA |
| 0345-G138-mut2 | G138-5/Mut2-3 | NIID0345 genomic DNA |
| 0345-G138-N248 | G138-5/N248-3 | NIID0345 genomic DNA |
| 0345-N248-mut2 | N248-3/Mut2-3 | NIID0345 genomic DNA |
| 0345-C | Mut2-5/LFH6 | NIID0345 genomic DNA |
| 0345-B | LFH3/LFH4 | NIID0345 genomic DNA |
| 35-A | LFH1/Mut1-3 | AfS35 genomic DNA |
| 35-mut1-mut2 | Mut1-5/Mut2-3 | AfS35 genomic DNA |
| 35-mut1-S138 | Mut1-5/S138-3 | AfS35 genomic DNA |
| 35-S138-mut2 | S138-5/Mut2-3 | AfS35 genomic DNA |
| 35-C | Mut2-5/LFH6 | AfS35 genomic DNA |
| 35-B | LFH3/LFH4 | AfS35 genomic DNA |
| Hph | Hph5/Hph3 | pHph plasmid DNA |

**Table 4.**
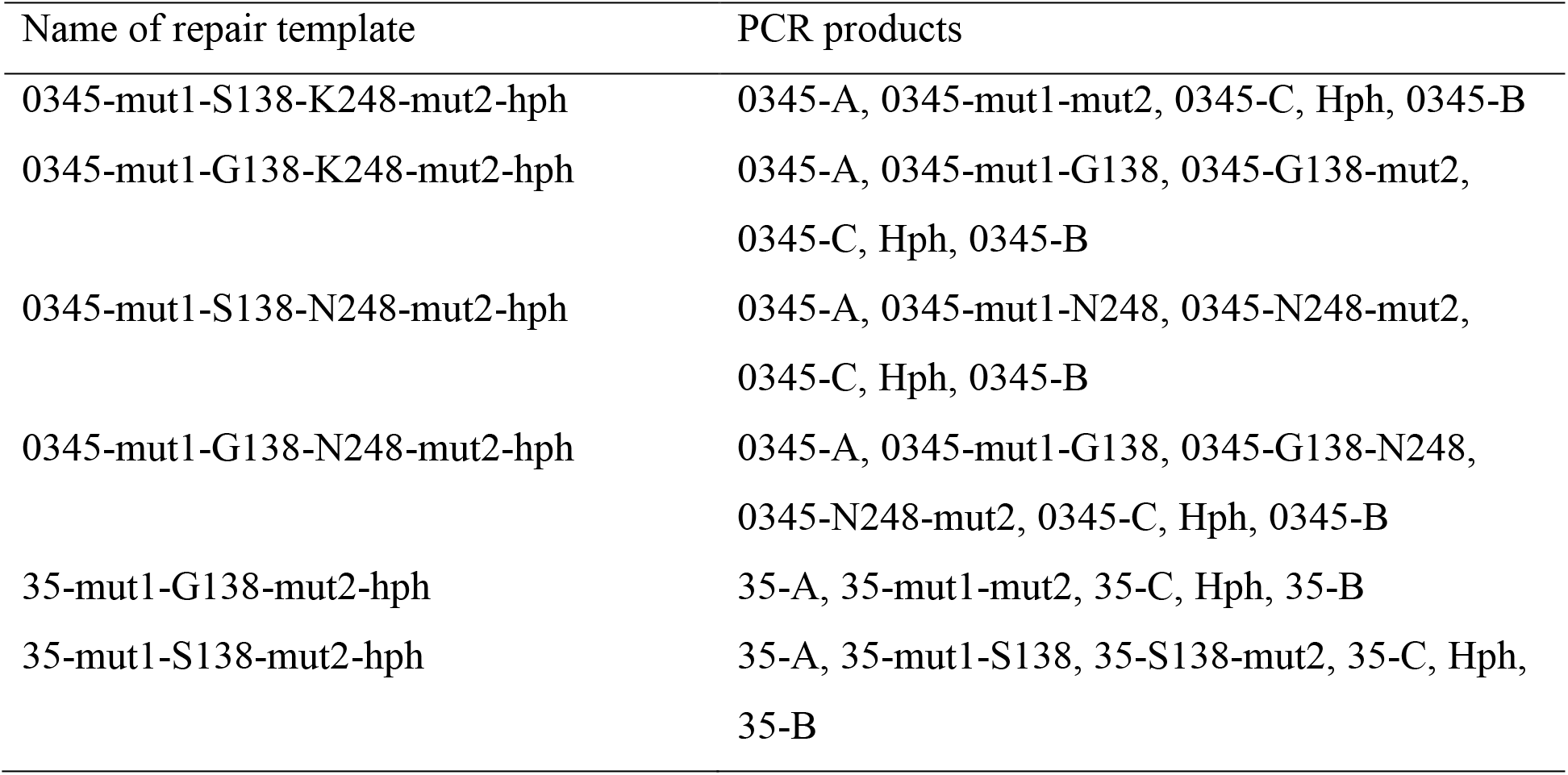
Combination of PCR products for repair template amplification used in this study.

### *A. fumigatus* transformation

*A. fumigatus* protoplasts were generated and fungal transformation was performed as previously described (13), with slight modifications. Briefly, conidia were incubated in YG medium for 6 h at 37°C. Following incubation, the cell walls of germlings were digested with 0.2 g/mL VinoTaste Pro (Novozymes, Denmark) for 1 h at 30°C; 20 pmol Cas9-NLS protein (New England Biolabs) and 10 pmol each *in vitro*-synthesized sgRNA1 and sgRNA2 were mixed and incubated for 25 min, generating ribonucleoproteins (RNPs). Protoplasts were transformed with 2–3 μg of repair templates and RNPs and plated onto CD supplemented with 1 M sucrose. Using NIID0345 clinical isolate as a host, repair templates 0345-mut1-S138-K248-mut2-hph, 0345-mut1-G138-K248-mut2-hph, 0345-mut1-S138-N248-mut2-hph, or 0345-mut1-G138-N248-mut2-hph were used to generate strains NIID0345-mut1-2, NIID0345-S138G, NIID0345-K248N, or NIID0345-S138G-K248N, respectively. Using AfS35 strain as a host, repair templates 35-mut1-G138-mut2-hph or 35-mut1-S138-mut2-hph were used to generate strains AfS35-mut1-2 or AfS35-G138S, respectively. Following a 15-h incubation at 37°C, plates were overlaid with CD top agar containing 400 μg/mL hygromycin. Positive colonies were confirmed by colony PCR using KOD FX Neo DNA polymerase (TOYOBO, Japan) with the primers Discheck5 and Discheck3 (which were designed at the region outside the repair template sequence), followed by nucleotide sequencing of *cyp51A*, including the promoter region.

### Antifungal susceptibility testing

Susceptibility to VRC, ITC, and POS were evaluated with Etest strips according to the manufacturer’s instruction (Biomerieux, France). Strains were grown at 37°C, and growth inhibition was visually evaluated after 48 h. Susceptibility tests were performed in three independent Cyp51A-sequence-confirmed transformants for each strain.

## Results

### Gly138Ser and Asn248Lys were found in the Cyp51A sequence of an azole-resistant clinical isolate

The susceptibilities of the clinical isolate from a patient with CPPA were determined by Etest methods. The isolate was not susceptible to VRC, ITC, or POS (Fig. 1), but was susceptible to amphotericin B, micafungin, and caspofungin (data not shown). We identified NIID0345 as an *A. fumigatus* strain via sequence analysis of the ITS and D1/D2 regions and the β-tubulin gene. Comparison of *cyp51A* from the azole-resistant isolate (NIID0345) with those from azole-susceptible strains (Af293 and AfS35) revealed that NIID0345 carried two amino acid substitutions: Gly138Ser (GGC→AGC) and Asn248Lys (AAT→AAA). These results indicated that one or both of these SNPs may be responsible for azole resistance.

**Figure 1.**
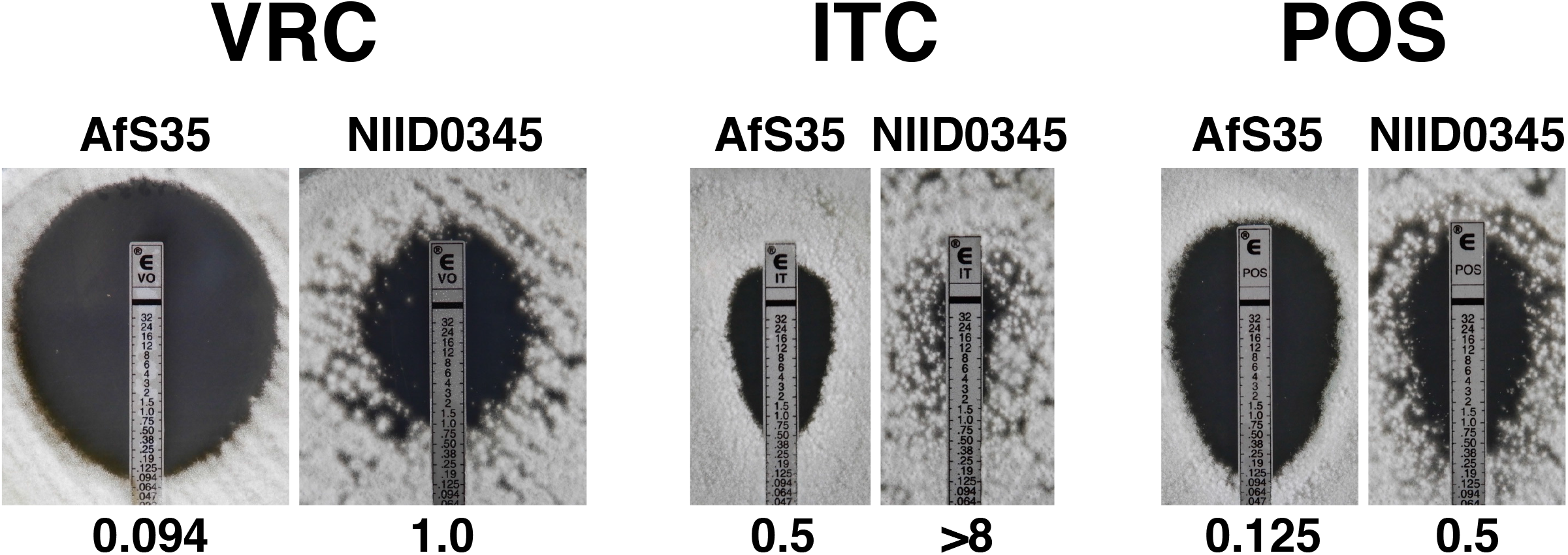
Antifungal susceptibility testing using Etest strips for voriconazole, itraconazole, and posaconazole in the azole-susceptible *Aspergillus fumigatus* strain AfS35 and clinical azole-resistant *A. fumigatus* strain NIID0345. The number below each photo represents the MIC (μg/mL).

### Cas9/CRISPR-mediated substitution of Serine at 138 to Glycine in *cyp51A* of NIID0345

To verify which SNP is involved in azole resistance, we substituted the nucleotide sequences corresponding to amino acid Ser138 and/or Lys248 in the clinical isolate NIID0345. We attempted to replace the genomic *cyp51A* gene locus with a linear DNA fragment harboring mutations by homologous recombination. It is well known that the rate of homologous recombination is very low in a host strain having an NHEJ repair pathway, such as clinical isolates. In addition, the Cas9/CRISPR system can be used to increase the efficiency of homologous recombination in *Candida glabrata* (15). Therefore, we used Cas9/CRISPR to create double-stranded breaks close to the N- and C-terminus of the *cyp51A* coding region, and to facilitate the replacement of the genomic *cyp51A* locus with a repair template DNA fragment via homologous recombination. Off-target effects are commonly encountered when Cas9 and sgRNA are continuously expressed via the introduction of plasmid or DNA; however, recent studies have shown that off-target effects were reduced by the direct introduction of the Cas9 protein and synthesized gRNA (16). Successful genome editing via the introduction of Cas9-gRNA ribonucleoprotein has also been reported in filamentous fungi, such as *A. fumigatus*. In the repair template DNA fragment, the nucleotide sequence responsible for amino acid substitutions thought to confer azole resistance in NIID0345 was substituted for a nucleotide sequence corresponding to that present in the azole-sensitive isolates. Furthermore, the construct was made Cas9/CRISPR resistant by introducing nuclease-resistant silent mutations into the two *cyp51A* gRNA target sites (Fig. 2). Cas9/sgRNA ribonucleoproteins and the repair template were simultaneously transformed via the protoplast-polyethylene glycol method into the *A. fumigatus* azole-resistant clinical isolate, and the transformants were selected with hygromycin. By confirming the nucleotide sequence of colony-directed PCR fragments of *cyp51A*, four kinds of recombinant strains were produced: a strain with only the nuclease-resistant mutation, those with Ser138Gly substitution, those with Lys248Asn substitution, and those with both Ser138Gly and Lys248Asn substitutions.

**Figure 2.**
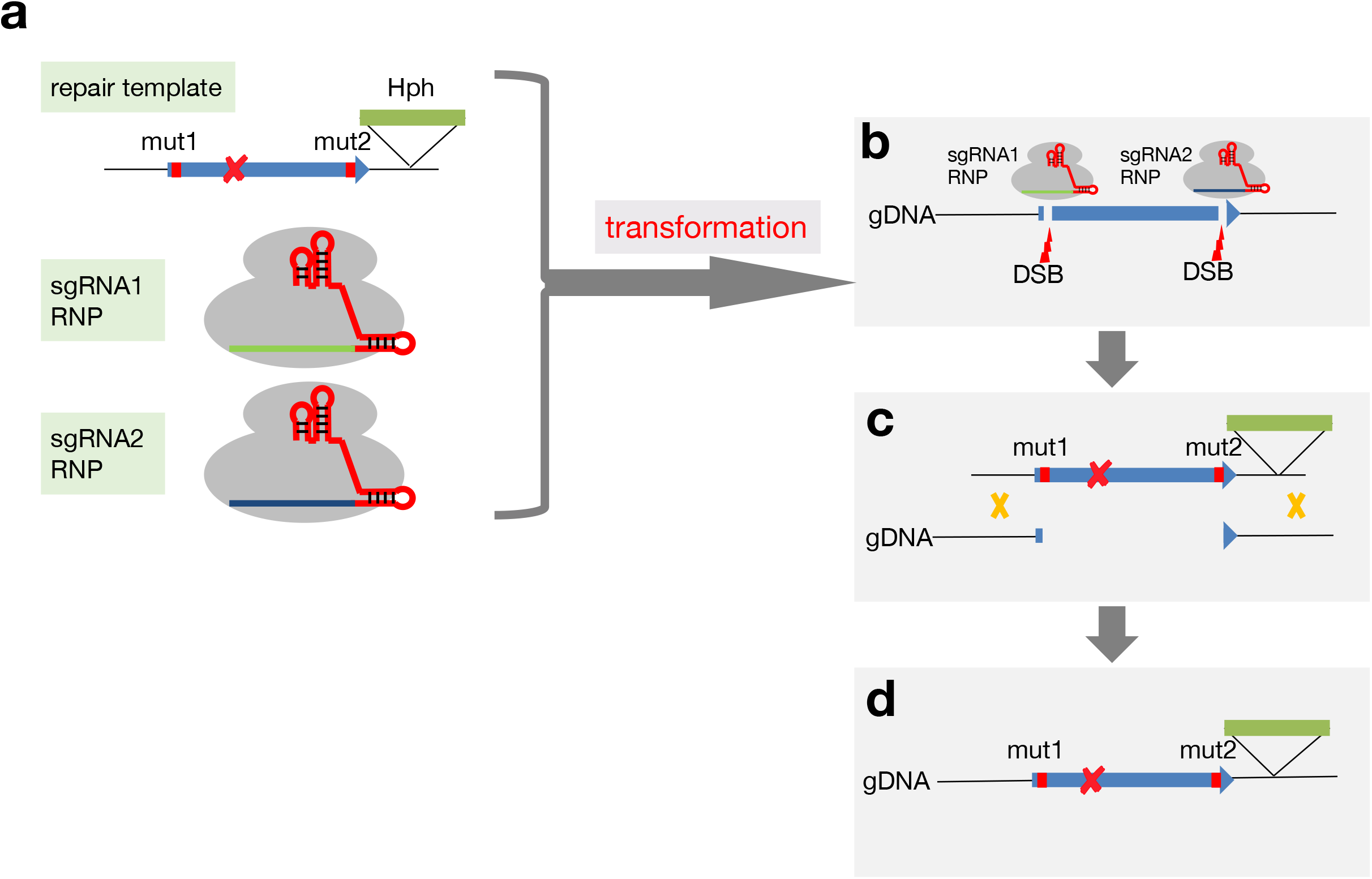
Overview of genetic modification via Cas9/CRISPR-promoted homology-directed repair. (a) Cas9 protein and *in vitro*-synthesized sgRNAs were mixed to form two RNPs. The repair template and two RNPs were transformed into *Aspergillus fumigatus* protoplasts. (b) The dual Cas9–sgRNA complex introduced two double-stranded breaks at the N-terminus and C-terminus of *cyp51A* (c and d). The cleaved *cyp51A* on the genomic DNA is replaced by the repair template, resulting in the introduction of the desired mutations and *hph* marker. The silent mutations mut1 and mut2 on the repair template and the replaced genomic DNA cannot be cleaved by RNPs nuclease.

Next, we examined azole susceptibility testing using Etest strips on the constructed recombinant strains. The strains in which only nuclease-resistant silent mutations were introduced demonstrated a similar azole-resistant profile as the parental strain NIID0345 (Fig. 3A), indicating that Cas9/CRISPR-mediated homologous recombination had no effect on azole susceptibility. Both of the recombinant strains with Ser138Gly and Ser138Gly/Lys248Asn amino acid substitutions showed increased susceptibilities to all azoles tested, whereas the strain with only Lys248Asn substitution showed a similar azole-resistant profile to the parental clinical isolate. These results indicate that Lys248 is not associated with azole resistance, and Ser138 is responsible for azole resistance in this clinical isolate.

**Figure 3.**
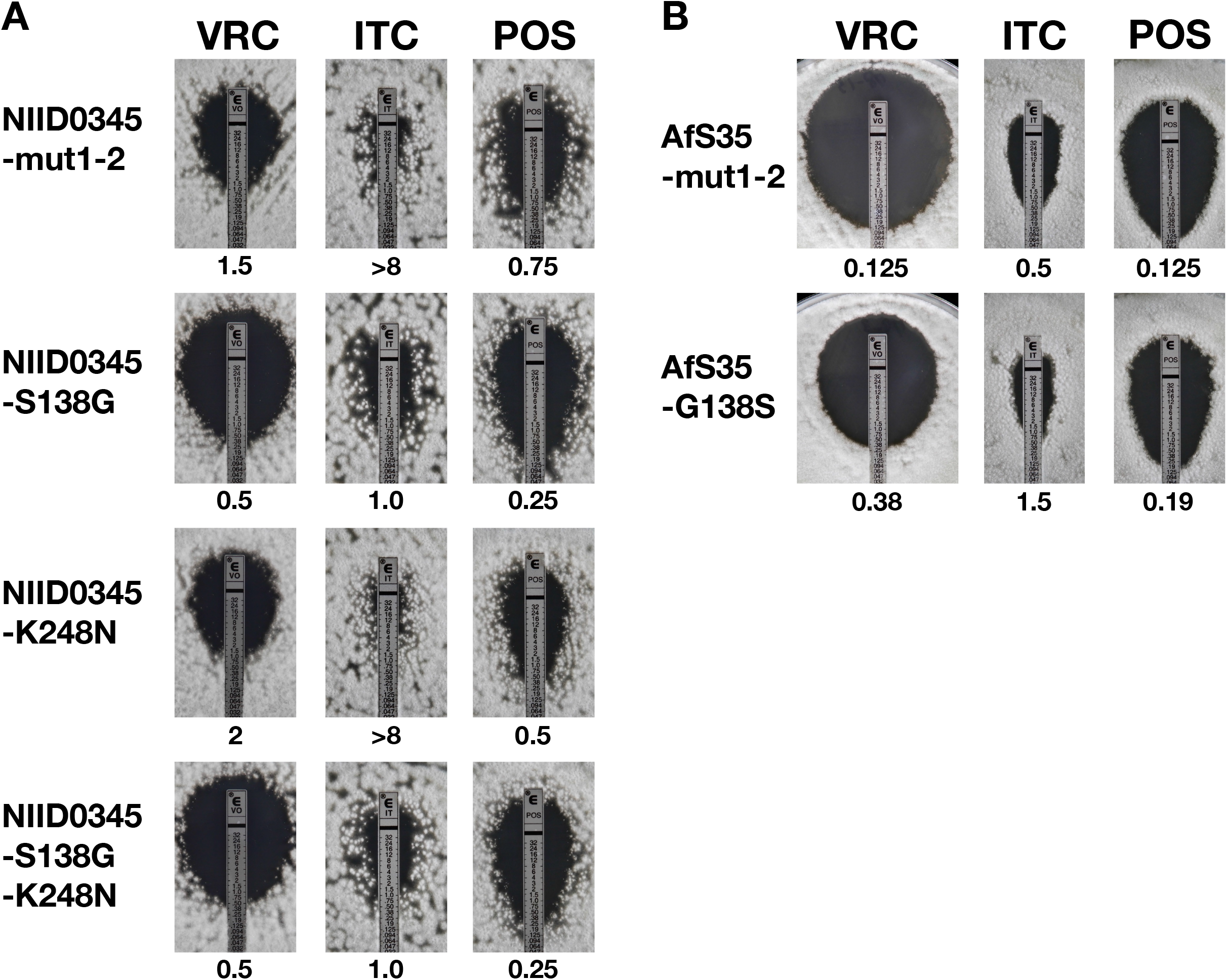
Antifungal susceptibility testing using Etest strips for voriconazole, itraconazole, and posaconazole for the strains generated via Cas9/CRISPR-promoted gene replacement from the strains NIID0345 (A) and AfS35 (B). The number below each photo represents the MIC (μg/mL).

### Gly 138 was substituted in the azole-susceptible strain AfS35

To verify whether Gly138 in Cyp51A is responsible for azole resistance, amino acid substitution of Gly138 to serine was introduced into the azole-susceptible strain AfS35. The method to produce the recombinant strain was the same as above; however, highly efficient homologous recombination was expected because the strain AfS35 is deficient in the NHEJ repair system. As expected, almost all transformants exhibited ideal recombination. We constructed two kinds of recombinant strains: a strain with only nuclease-resistant silent mutations and one in which Gly138 was substituted with Ser138. Azole susceptibility testing by Etest of the recombinant strains showed a slight decrease in azole susceptibility when the Ser138 mutation was introduced into the azole-susceptible strain AfS35 (Fig. 3B). The strain with only nuclease-resistant silent mutations demonstrated a similar azole susceptibility profile as the parental strain AfS35. From these results, we elucidated the direct involvement of Gly138 in Cyp51A in azole resistance, which had previously been supported only by indirect epidemiological evidence.

## Discussion

The Cas9/CRISPR genome editing technique used in this study has enabled site-directed mutagenesis, altering Ser138 to glycine on the genomic Cyp51A locus in an azole-resistant clinical strain. This is, to our knowledge, the first example of site-directed mutagenesis performed in a clinical, azole-resistant fungal isolate to elucidate whether azole susceptibility is altered by mutations in the genomic Cyp51A locus. Although many azole-resistant isolates with SNPs in *cyp51A* have been identified, there have been few studies that genetically confirm their correspondence to low azole susceptibility. Although genetically recombinant strains harboring mutations such as TR34-Leu98His (17), TR46-Tyr121Phe-Thr289Ala (18), Gly54Trp (17), and Thr301Ile (19) in the genomic Cyp51A locus have been reported to date, all these strains were constructed in the *akuB*/Ku80-deficient strain as a recipient. It is well known that wild-type strains, such as clinical isolates, tend to exhibit low efficiency in homologous recombination, largely because of high NHEJ activity. To overcome this limitation, the gene encoding either KU70 or Ku80, which are the components of NHEJ machinery, was knocked out, leading to a significant increase in the frequency of homologous recombination (14, 20). In contrast, our method using Cas9/CRISPR can facilitate efficient homologous recombination without the inactivation of the NHEJ pathway, which is supported by previous studies, concluding that the frequency of homologous recombination can be increased by the Cas9/CRISPR system in *C. glabrata* (15).

To build the Cas9/CRISPR system in *A. fumigatus* clinical isolates, we incorporated several additional methods to improve the efficiency and accuracy of *cyp51A* gene replacement events. Improved efficiency of *cyp51A* replacement was achieved by introducing two DSBs via the design of two target sequences for sgRNA at sites close to the N-terminus and C-terminus, repressing homologous recombination within the *cyp51A* coding region. Additionally, to avoid digestion of the repair template and re-digestion of the edited target after the homologous recombination event, nuclease-resistant silent mutations were introduced in two loci of three codons immediately upstream from the PAM sites of the repair template, preventing it from being targeted by Cas9/CRISPR(21). To minimize off-target effects from DNA-based continuous Cas9 and sgRNA expression (which should be considered whenever Cas9/CRISPR system is used for genome editing), we introduced ribonucleoproteins consisting of commercially available recombinant Cas9 protein and *in vitro*-synthesized sgRNAs directly into protoplasts of clinical isolates. As one means of minimizing off-target effects, directly transfected Cas9 protein reduces the off-target cleavage rate when compared with Cas9 expression by a plasmid or mRNA transfection in mammalian cells (16). One recent study has demonstrated that direct delivery of Cas9–gRNA ribonucleoprotein can facilitate genome editing in *A. fumigatus* (10). Based on these improvements, we produced a simple, efficient, and accurate site-directed mutagenesis system to investigate structure–phenotype relationships of the azole target Cyp51A. Since this system can be applied to numerous genes other than *cyp51A*, this method will accelerate the progress of many pathogenic fungal studies.

Multiple Gly138 polymorphisms in *cyp51A* have been identified in *A. fumigatus* azole-resistant isolates, most of which alter Gly/GGC to Cys/TGC and Ser/AGC (22–25). Although *Saccharomyces cerevisiae* expressing *cyp51A* with the Gly138Cys mutation showed reduced susceptibility to all three azoles compared with the control strain, no genetic studies using *A. fumigatus* as a host have been performed to date to investigate the function of Gly138 in Cyp51A. We isolated an azole-resistant *A. fumigatus* strain from a CPPA patient with long-term VRC treatment and identified two amino acid substitutions—Gly138Ser and Asn248Lys—which, when compared with Cyp51A nucleotide sequences from azole-susceptible strains, may be potential polymorphisms conferring azole resistance. According to the homology model structure of the Cyp51A protein, Gly138 is located in a channel 1 helix close to the heme cofactor, and a mutation at this position could disturb the heme environment, which may lead to multiple azole resistance (26, 27). Structure modeling and epidemiology have predicted that the Gly138Ser mutation is the amino acid responsible for azole resistance, and our Cas9/CRISPR gene replacement system has molecularly confirmed that the Ser138 is responsible for the pan-azole-resistant phenotype in the clinical isolate NIID0345. However, genome-edited mutants harboring Gly138 in the NIID0345 genetic background have exhibited much lower susceptibility to all azoles tested than the strain AfS35, in which the amino acid at position 138 is intrinsically glycine. Similarly, the genome-edited mutant harboring Ser138 in AfS35 genetic background has exhibited higher susceptibility than the strain NIID0345, in which the amino acid at position 138 is intrinsically serine. These results indicate that the strain NIID0345 may have low azole susceptibility for reasons other than alteration in *cyp51A*. Additional genomic analysis is needed to identify unknown, non-Cyp51A mechanisms of azole resistance in the clinical isolate NIID0345.

In conclusion, we have developed a simple, efficient, and accurate gene replacement system using Cas9/CRISPR genome editing techniques, and applied these techniques to investigate the mechanisms of azole resistance via Cyp51A alteration. We confirm at the molecular level that the Gly138Ser mutation is one reason for azole resistance in a clinical isolate. There are many *cyp51A* mutations that may result in potential, but unconfirmed amino acid changes conferring azole resistance. Further investigation of Cyp51A using our Cas9/CRISPR system is required to verify whether the diverse SNPs reported to date are in fact responsible for azole resistance.

## Acknowledgment

We thank Hiroko Tomuro for technical assistance. This work was supported by MEXT KAKENHI Grant Number JP126K09954 and Joint Usage/Research Program of Medical Mycology Research Center, Chiba University (17–9). The authors would like to thank Enago (www.enago.jp) for the English language review.

We declare the authors have no conflicts of interest.

